# Occurrence, antimicrobial resistance pattern and molecular characterization of *Listeria monocytogenes* isolated from bovine’s milk and meat in Mekelle City, Ethiopia

**DOI:** 10.1101/2021.11.16.468824

**Authors:** Tesfay Hailu Kidanu, Getachew Gugsa, Yisehak Tsegaye Redda, Meselu Ahmed, Nesibu Awol

**Author notes:** Corresponding author: Getachew Gugsa E-mail. These authors contributed equally to this work.

## Abstract

*Listeria monocytogenes* is an opportunistic and emerging foodborne zoonotic pathogen that encompasses a diversity of strains with varied virulence and can cause serious human and animal infections worldwide with low incidence but high hospitalization and case fatality rates. A cross-sectional study was conducted from December 2016 to June 2017 to estimate the molecular epidemiology of *L. monocytogenes* and its serotypes, and antimicrobial resistance pattern of isolates in Mekelle City. A total of 768 (384 of milk and 384 meat) samples of bovine origin were collected using a purposive random sampling technique. Isolation and identification of *L. monocytogenes* was done according to standard and recommended bacteriological procedures. Genome-based confirmation of each isolate was performed at species and serovar levels by targeting *Iap, Imo0737, ORF2819* and *ORF2110 genes* using specific primers. *In vitro* antimicrobial susceptibility testing was performed using agar plate antibiotic disk diffusion method. The overall prevalence of *L. monocytogenes* was 26 (3.39%). Sample type prevalence rates of *L. monocytogenes* were 4.17 % and 2.6% in meat and milk samples, respectively. There was a statically significant difference (p<0.05) on the prevalence rates of the organism in meat samples collected from abattoir (1.67%), butcher shops (8.33%), and restaurants (8.33%). Serovars that were identified were belonged to 1/2b and 4b. Large proportions of isolates were highly susceptible to Ampicillin (88.46%) and Vancomycin (84.62%). However, the isolates had shown the highest level of resistance against Nalidixic Acid (96.15%). The highest intermediate was observed to Amoxicillin (57.69%). Moreover, 42.31% of the isolates were developed resistance for more than two drugs. Hence, both its occurrence and development of a multi-drug resistance indicated that, a coordinated effort is imperative to reduce or eliminate the risk posed by this pathogen in food chains and on controlled and careful use of antimicrobials both in veterinary and human treatment regimes.

## Introduction

Food-borne diseases remain a major public health problem across the globe. The problem is severe in developing countries due to difficulties in securing optimal hygienic food handling practices and with the increase in consumption of raw products of animal origin. Hence, food safety and particularly the safety of products of animal origin is an increasingly significant issue concerning human health [1-3]. *Campylobacter species* (*Campylobacter* spp.), *Salmonella spp*., *Listeria spp*., and *Escherichia coli* O157:H7 have been generally found to be responsible for majority of food-borne outbreaks [4, 5]. Out of the agents associated with the current worldwide increase in cases of food-borne diseases, *L. monocytogenes* is a concern especially in the developing world [6].

*Listeria* are Gram positive, non-spore forming, non-capsular, short, regular rods, motile with a characteristic tumbling movement, and with a low G+C content. The six species of *Listeria* are *L. monocytogenes, L. ivanovii, L. seeligeri, L. innocua, L. welshimeri*, and *L. grayi*. However, only *L. monocytogenes* and *L. ivanovii* are considered virulent. Human cases of *L. ivanovii* infection are rare [7-9], whereas *L. monocytogenes* is an important food-borne pathogen [10]. *L. monocytogenes* is the causative agent of listeriosis, a food-borne disease that affects mainly at risk populations such as the elderly, pregnant woman, newborns and immunocompromised individuals [11].

*L. monocytogenes* comprises 13 serotypes with different virulence potential namely: 1/2a, 1/2b, 1/2c, 3a, 3b, 3c, 4a, 4ab, 4b, 4c, 4d, 4e, and 7 based on somatic (O factor) and flagella (H factor) antigens [12, 13]. However, more than 95% of human listeriosis commonly associated with isolates that belong to serotypes 1/2a, 1/2b, and 4b, and of these serotype 4b has been related to the most recent outbreaks of listeriosis, and serotypes 1/2a and 4b are commonly reported in animals [14-20].

*L. monocytogenes* can be found in a wide variety of raw and processed foods including dairy products, raw meat, vegetables, and fishery products foods of animal origin. Milk and dairy products, and various meats and meat products have all been associated with *Listeria* contamination and with outbreaks or sporadic cases of listeriosis [21-24, 12]. *L. monocytogenes* is transmitted from man to man or from animal to man mostly through ingestion of the organism with contaminated raw milk, raw meat, and dairy products. Hence, these food products have been connected in several outbreaks of listeriosis [25]. The central characteristics of *L. monocytogenes* contributing to food-borne transmission are the ability to grow as low as -0.4°C, resist heat, salt, nitrite, acidity, endure osmotic stress, and survive mild preservation treatment measures commonly used to control the growth of organisms in food [26]. However, in most African countries, there are a few reports on *Listeria* and listeriosis, as compared to the Europe and USA [27].

Despite the poor surveillance programmes and lack of epidemiological data to establish a comprehensive incidence rate of human listeriosis in Africa, there are a number of studies that reported the occurrence of *L. monocytogenes* in foods from African countries [28-31].

*L. monocytogenes*, as well as other *Listeria* spp., are usually susceptible to a wide range of antibiotics [32]. However, *L. monocytogenes* strains resistant to one or more antibiotics have been recovered from food, the environment and sporadic cases of human listeriosis [33-35]. Moreover, various studies have reported *L. monocytogenes* strains that showed multiple drug resistance to different antimicrobial substances, which is significance for the public health, therefore requiring an arduous monitoring of the antimicrobial susceptibility of *L. monocytogenes* strains [36, 37]. In Ethiopia, the actual epidemiological data about the prevalence as well as antimicrobial susceptibility pattern of *L. monocytogenes* is lacking both in the veterinary and public health sectors. Moreover, there is no published research work done on the epidemiology and antimicrobial susceptibility patterns of *L. monocytogenes* and its strain in foods of bovine origin, particularly in raw milk and meat in the current study area. Hence, this study was stipulated to estimate the molecular epidemiology of *L. monocytogenes* and its serotypes, and determine the antimicrobial susceptibility of *L. monocytogenes* from raw milk and raw meat samples of bovine origin collected from different sources in Mekelle City, Ethiopia.

## Materials and Methods

### Ethics approval and consent to participate

This study was reviewed and approved by the Research Ethics Committee of the College of Veterinary Sciences, Mekelle University. Owners, managers and workers of the different sites were informed on the procedures and significance of the study. Each data and analysis result was kept confidential and communicated to concerned bodies. Any participants who were not volunteers were not forced to be included.

### Study area

The study was conducted from December 2016 to June 2017 in Mekelle City. Mekelle is the capital city of Tigray Regional State, which is located at 39° 29`E and 13°-30` N at an average altitude of 2000 m. a. s. l. and 783Km far from Addis Ababa. The climate of the study area conforms to that of Ethiopian Highlands. The mean annual rainfall is 619mm, and is bimodal with a short rainy season occurring from March to May and another from middle September to February. The annual minimum and maximum temperature is 11.8°C and 29.9°C, respectively [38]. It is administratively divided into 7 sub cities and is subdivided to 33 Kebelles.

### Study design and study population

A cross-sectional study was conducted from December 2016 to June 2017 in Mekelle City. The study populations comprised of purposively selected milking dairy cows and slaughtered cattle found in Mekelle City. The samples were collected from randomly selected dairy farms, milk-shops, and cafeteria for bovine milk; and abattoir, butcher shops, and restaurants for beef. The samples were examined to estimate the prevalence of *L. monocytogenes* and determine its antimicrobial sensitivity pattern following the standard protocols and the species and serovars were further confirmed using species-specific and serovar specific PCR primers.

### Sampling size and sample collection

A total of 768 samples of bovine origin, comprised of milk (n=384) and meat (n=384), were collected using a purposive random sampling technique. The samples were collected based on the willingness of the owners until the required sample size was achieved. Thirty (30) ml of bovine milk sample was collected from dairy farms (166), shops (120), and cafeteria (98). At the same time, 25gm of bovine meat samples were collected from abattoir (240), butcher shops (84), and restaurants (60) using sterile plastic bags. All samples were labeled and transported with icebox to Molecular Biotechnology Laboratory of College of Veterinary Sciences, Mekelle University at the date of collection and stored at 4°C till further analysis was done.

### Bacteriological isolation

The procedure described by the International Organization for Standardization [39] was used for milk samples. While the techniques recommended by the International Standards Organization [40, 41] and the French Association for Standardization [42] with minor modification were employed for the isolation and identification of *L. monocytogenes* from meat.

#### Selective enrichment

The raw milk sample was thoroughly homogenized and inoculated into Listeria Enrichment Broth (LEB) at 1:9 ratios [43]. After 48hrs of incubation at 30°C, the sample from LEB suspensions was inoculated into Secondary Fraser Enrichment Broth (SFEB) and incubated at 35^º^C for 24 hrs. Similarly, for the meat samples, 1:9 proportion of sample to LEB was placed in sterile Stomacher plastic bags and homogenized for 1 min in laboratory stomacher (Lab Belender 400, Seward Medical, London, England). Then, the resulting suspension was incubated for 48 hrs. After 48hrs of incubation, 0.1ml of the culture was transferred to previously prepared SFEB and then incubated for 24hrs.

#### Selective plating

After 24hrs of incubation, aliquots of positive SFEB medium with black/dark brown or dark green color was taken and streaked onto Listeria Oxford Agar plates containing selective media with manufacturer’s supplements and the plates were incubated at 35°C for 24-48 hrs. Then, the growth of *Listeria species* on Listeria Oxford Agar plate was examined.

#### Confirmatory tests

Suspected *Listeria* colonies were sub-cultured onto Tryptose Soya Agar plate (TSA) and incubated at 37°C for 18-24 hrs. Those suspected *Listeria* colonies were characterized by using Gram’s staining, oxidase test, characteristics of hemolysis [43], carbohydrate utilization (xylose, rhamnose), and CAMP (Christie Atkins Munch Peterson) test following standard methods [41-43].

### Genomic DNA extraction and PCR amplification of the isolates

#### Genomic DNA extraction

The DNA extraction was performed using the phenol-chloroform method for all the isolates from samples of milk and meat according to [44].

#### The PCR conditions

Following preliminary bacteriological identification and genomic DNA extraction of *L. monocytogenes* isolates, genome-based confirmation of each isolate was performed. The PCR condition that was used in this study has been explained previously by [45] and sequence of primers by [46] for *L. monocytogenes* and [47] for its serovars. As a positive control, a standard strain of *L. monocytogenes* was used. The primers used for the PCR reaction were F-3’-TTA TAC GCG ACC GAA GCC AAC-5’ and R-3’-CAA ACT GCT AAC ACA GCT ACT-5’ for amplification of *Iap gene* of *L. monocytogenes* (660 bp); F-5’-AGG GCT TCA AGG ACT TAC CC-3’ and R-5’-ACG ATT TCT GCT TGC CAT TC-3’ for amplification of *Imo0737 gene* of *L. monocytogenes* serovar 1/2a (691 bp); F-5’-AGC AAA ATG CCA AAA CTC GT-3’ and R-5’-CAT CAC TAA AGC CTC CCA TTG-3’ for amplification of *ORF2819 gene* of *L. monocytogenes* serovar1/2b (471 bp); F-5’-AGT GGA CAA TTG ATT GGT GAA-3’ and R-5’-CAT CCA TCC CTT ACT TTG GAC-3’ for amplification of *ORF2110 gene* of *L. monocytogenes* serovar 4b (597 bp).

Amplification was carried out with thermal cycling conditions of an initial denaturation at 95°C for 5 min followed by 35 cycles of denaturation at 95°C for 45s, annealing at 50°C for 1 min, and extension at 72°C for 1 min, and with a final extension at 72°C for 10 min for detection of *L. monocytogenes*. But the detection of serovars was optimized with cycling conditions of an initial denaturation at 94ºC for 5 minutes followed by 35 cycles of denaturation at 94ºC for 30s, annealing at 54ºC for 15s, and extension at 72º C for 75s, with final extension at 72ºC for 10 min. Finally, the PCR products along with DNA molecular weight marker (3B Black Bio Biotech) were separated by running on a 2% (w/v) agarose gel containing 0.3mg/ml ethidium bromide. Electrophoresis was conducted in a horizontal equipment system for 120 min at 90 V using 1X TAE buffer. The amplicons were visualized under a gel documentation system and their molecular weights were estimated.

### Antimicrobial susceptibility testing

All the 26 phenotypically and molecularly characterized isolates of *L. monocytogenes* were tested for antibiotic susceptibility pattern. The method applied for the *in vitro* antimicrobial susceptibility testing of *L. monocytogenes* isolates was the agar plate antibiotic disk diffusion method using Kirby-Bauer technique [48]. The following thirteen antibiotic disks (Himedia Laboratory Pvt limited, Mumbai, India) with their concentrations given in parentheses were used in the antibiogram testing: Amoxicillin (25μg), Ampicillin (10μg), Ciprofloxacillin (5μg), Clindamycin (10μg), Cloxacillin (5μg), Erythromycin (15μg), Gentamycin (10μg), Methicillin (30μg), Nalidixic Acid (30μg), Penicillin G (10μg), Streptomycin (10μg), Tetracycline (30μg), and Vancomycin (30μg). The selection of these antibiotics was based on the availability and frequent use of these antimicrobials in the study area both in veterinary and human medicine. The results were interpreted as Susceptible (S), Intermediate (I), and Resistant (R) categories based on the critical points recommended by the Clinical and Laboratory Standards Institute [49].

### Data management and processing

The data collected from the results of the laboratory investigation were entered into Microsoft Excel Spreadsheet (Microsoft Crop., Redmond, WA, USA) and transferred to STATA Version 11.1 for statistical analysis. Descriptive statistics, Chi-square test (χ^2^) and logistic regression were applied to see the associations of the food type and corresponding source as risk factors with that of the occurrence of the positive result; the degree of association was determined using Odds ratio (OR) and 95% confidence interval (CI) as explained by [50]. For those that test, a p-value less than 0.05 was considered statistically significant.

## Results

### Overall prevalence of *L. monocytogenes*

The overall prevalence of *L. monocytogenes*, from the total 768 samples, was found to be 26(3.39%). The sample type prevalence of *L. monocytogenes* was found to be 4.17 % and 2.6% in meat and milk samples, respectively. In general, the prevalence of *L. monocytogenes* was varied between sample types and sources. The prevalence of *L. monocytogenes* in meat samples collected from abattoir, butcher shops, and restaurants were 1.67%, 8.33%, and 8.33%, respectively. The result was significantly higher (p<0.05) in butcher shops and restaurants than abattoir. Whereas, out of 384 milk samples collected from dairy farms, milk shops, and cafeteria, the prevalence was 1.81%, 2.50%, and 4.08%, respectively. However, there was no statically significant difference (p>0.05) in the prevalence of *L. monocytogenes* among the various sources of milk samples (Table 1).

**Table 1.** Overall prevalence of *L. monocytogenes* from different sample sources.

| Sample type | Place of collection | No. Examined | Prevalence (%) | 95% CI | p-value |
| --- | --- | --- | --- | --- | --- |
| **Milk** | Farms | 166 | 3(1.81) | 0.0 - 1.00 | 0.531464 |
|  | Shops | 120 | 3(2.50) | 0.27 - 7.02 |  |
|  | Cafeterias | 98 | 4(4.08) | 0.51 - 10.55 |  |
|  | **Total** | **384** | **10 (2.6)** |  |  |
| **Meat** | Abattoir | 240 | 4(1.67) | 0.27 - 4.90 | 0.005659 |
|  | Butchers | 84 | 7(8.33)<sup>a</sup> | 1.24 - 19.62 |  |
|  | Restaurants | 60 | 5(8.33) | 0.84 - 17.87 |  |
|  | **Total** | **384** | **16 (4.17)** |  |  |
| **Grand Total** |  | **768** | **26(3.39)** |  | **0.0417** |

### PCR confirmation of *L. monocytogenes* isolates and serotyping

All the 26 phenotypically characterized *L. monocytogenes* isolates were also molecularly confirmed as *L. monocytogenes* using *L. monocytogenes* species-specific primer, which targets *Iap gene* as it is depicted below in Figure 2.

**Fig1.**
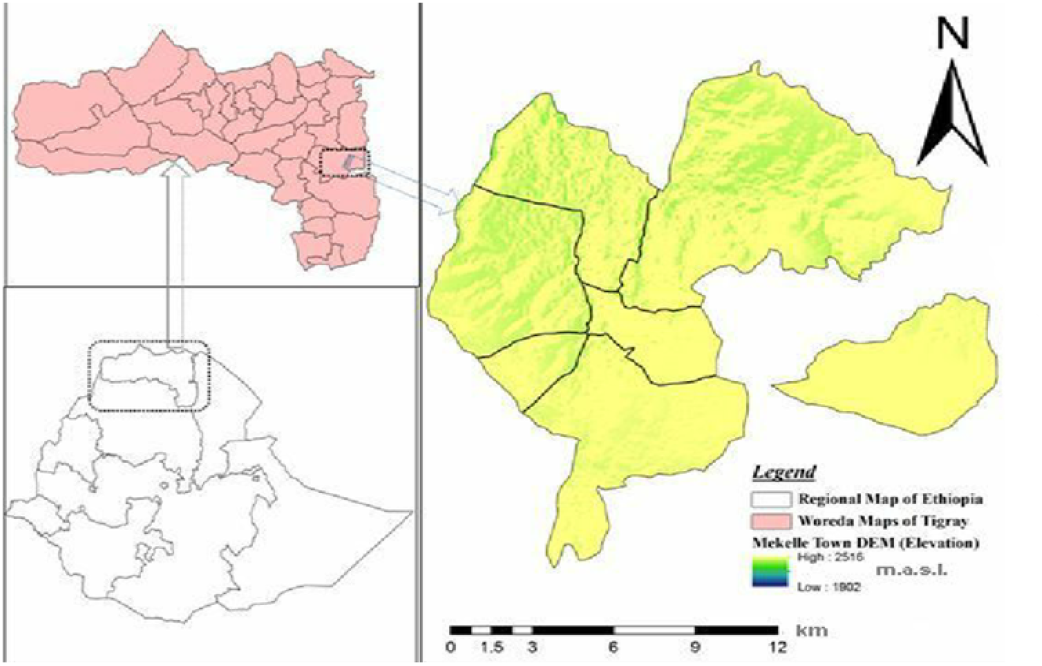
Map of the study area.

**Fig 2.**
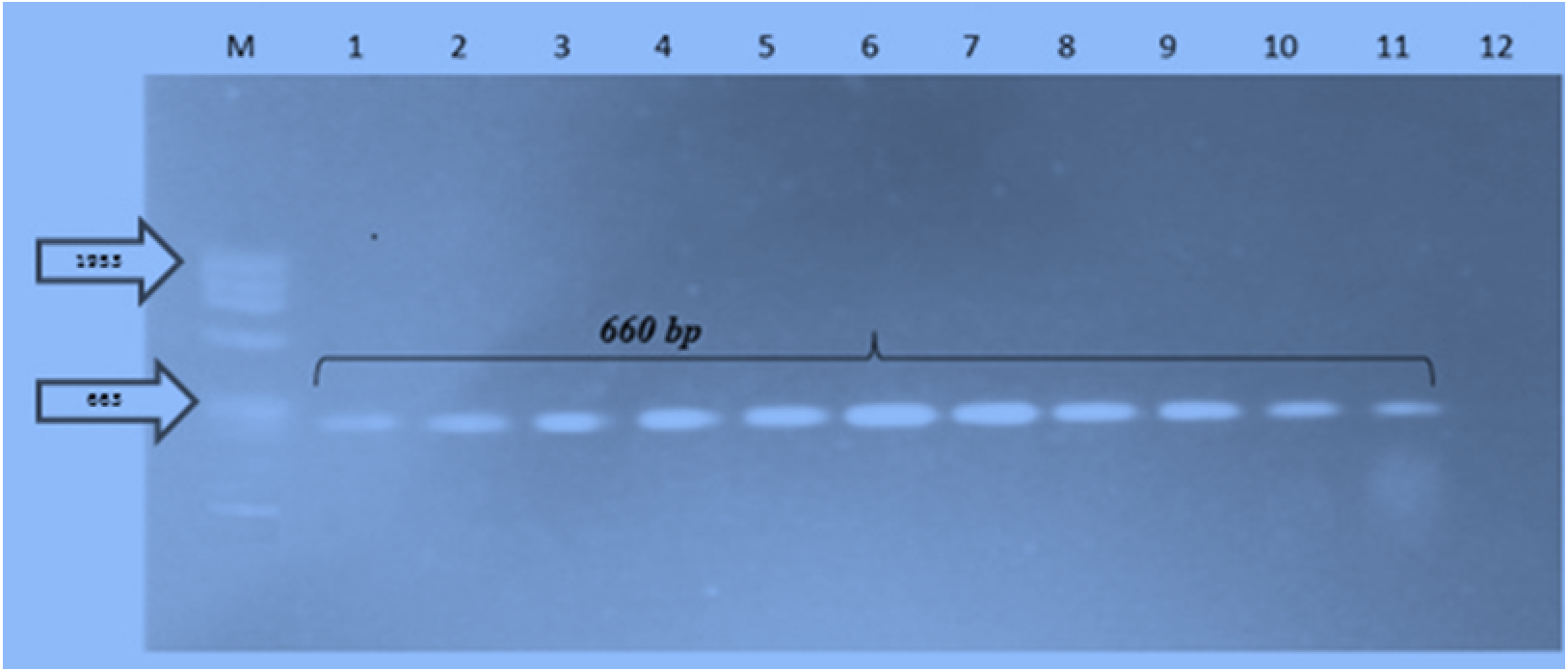
PCR Product of *L. monocytogenes*; M-DNA Ladder, Lanes 1-11-*L. monocytogenes* (660bp) and Lane 12-Negative control.

Moreover, the molecularly confirmed *L. monocytogenes* isolates were further identified at serovar levels using serovar specific primers with the help of multiplex PCR. The serovars that were identified were belonged to 1/2b and 4b as it shown below in Figure 3.

**Fig 3.**
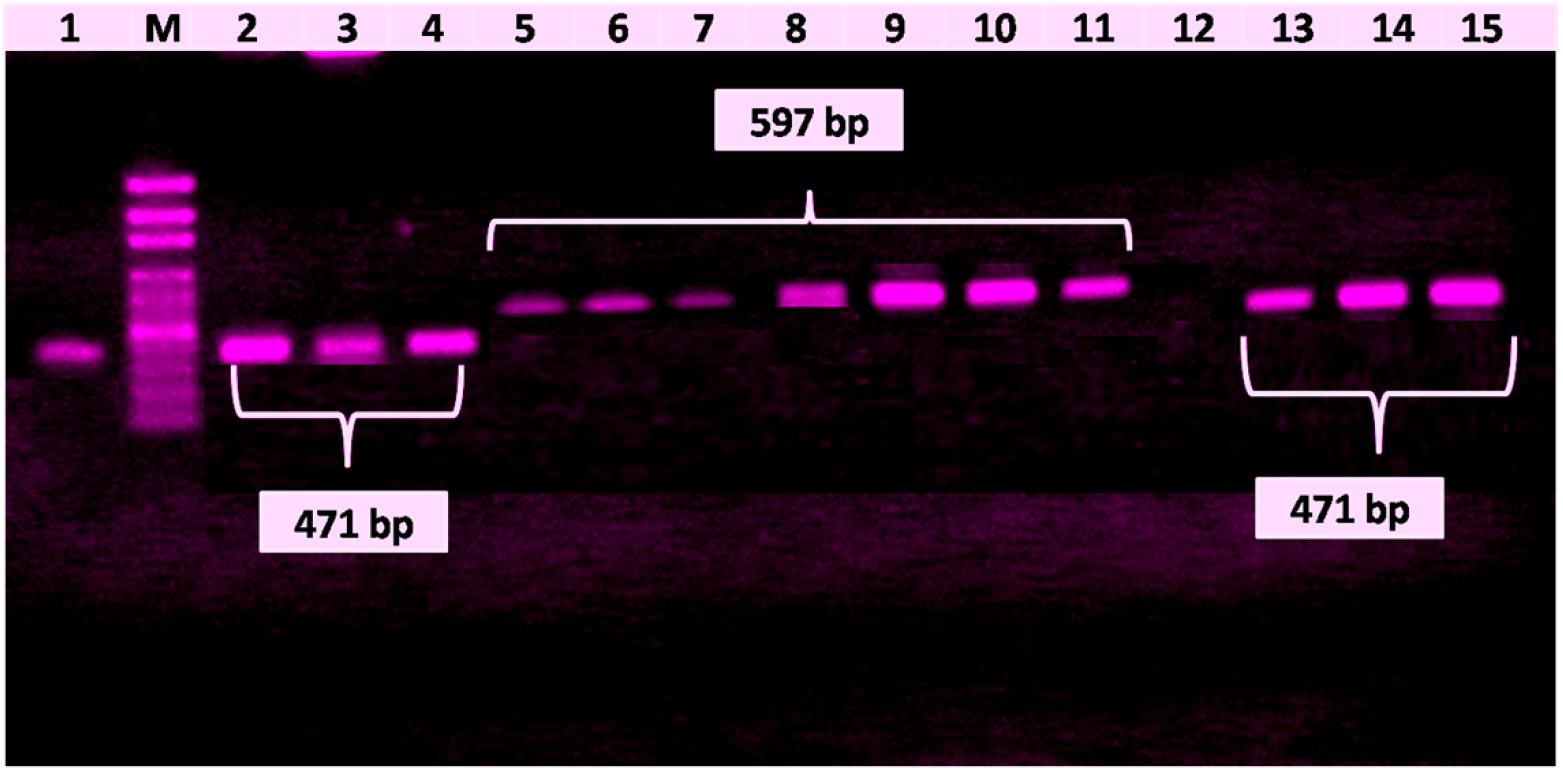
PCR Products: M-DNA ladder; Lanes 1-4 and Lanes 13-15 for serovar 1/2b(471bp); and Lanes 5-11 for serovar 4b (597 bp); and Lane12-Negative control.

**Fig 4.**
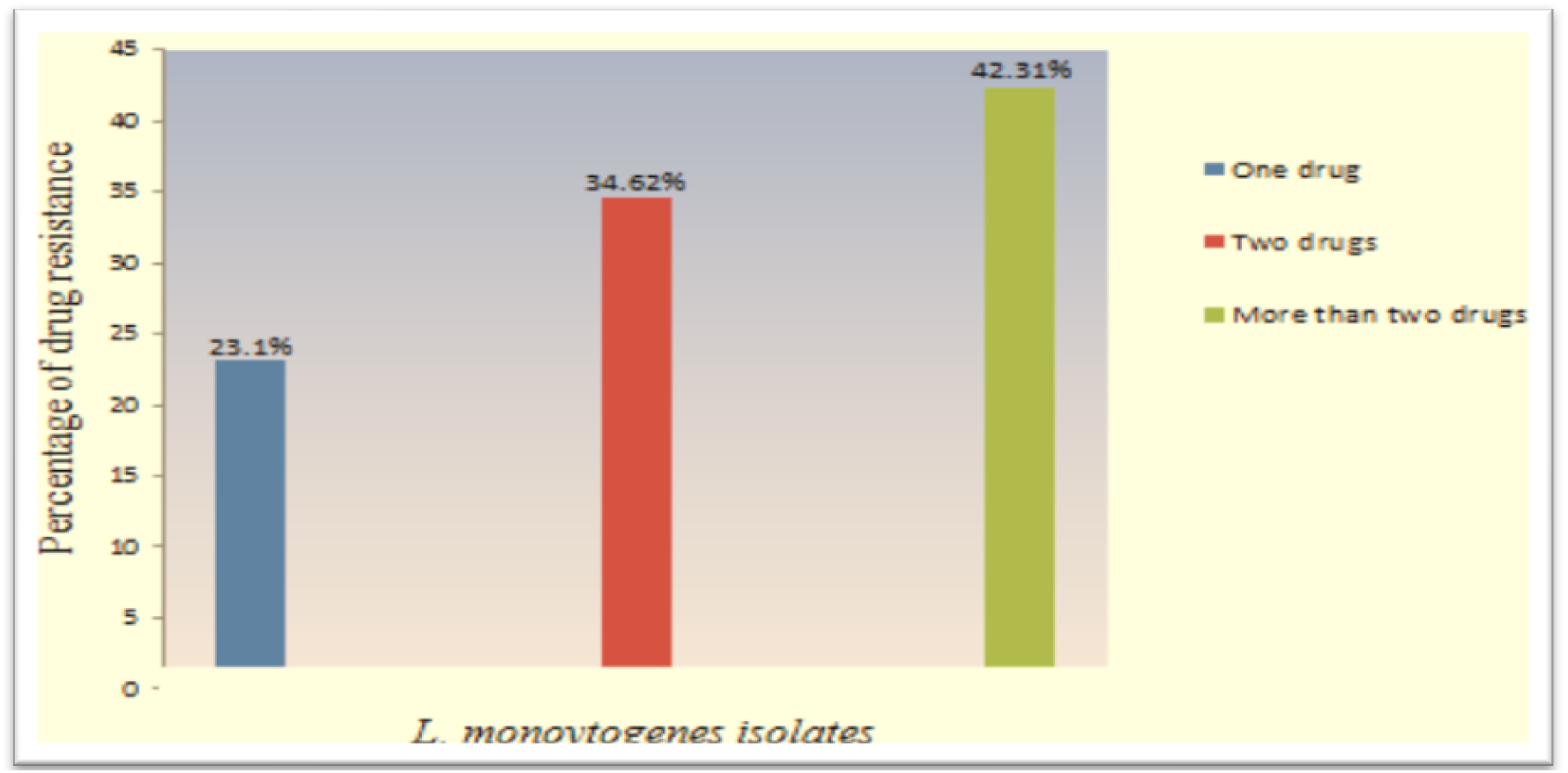
*In vitro* antimicrobial multidrug resistance pattern of *L. monocytogenes* isolates.

### Antimicrobial susceptibility pattern

All the 26 isolates of *L. monocytogenes* were tested for antibiotic susceptibility pattern. The antibiogram profile results indicated as large proportions of the isolates of this study were found to be highly susceptible to Ampicillin (88.46%) and Vancomycin (84.62%). However, the isolates had shown the highest level of resistance against Nalidixic Acid (96.15%). The highest intermediate was observed to Amoxicillin (57.69%) (Table 2). Moreover, 34.62%, and 42.31% of the isolates were developed resistance for two and more than two drugs, respectively, as it is shown in Figure 3.

**Table 2.** *In vitro* antimicrobial susceptibility pattern of *L. monocytogenes* isolates.

| Antimicrobials |  | Interpretations |  |
| --- | --- | --- | --- |
| Agents | No. of Susceptible (%) | No. of Intermediate (%) | No. of Resistance (%) |
| Penicillin G | 13(50.00) | 10(38.46) | 3(11.54) |
| Amoxicillin | 9(34.62) | 15(57.69) | 2(7.69) |
| Ampicillin | 23 (88.46) | 3(11.54) | 0(0) |
| Vancomycin | 22(84.62) | 4(15.38) | 0(0) |
| Ciprofloxacin | 20(76.92) | 6(23.08) | 0(0) |
| Streptomycin | 11(42.31) | 12(46.15) | 4(15.38) |
| Nalidixic Acid | 0(0) | 1(3.85) | 25(96.15) |
| Gentamycin | 21(80.77) | 5(19.23) | 0(0) |
| Methicillin | 19(73.07) | 7(26.92) | 0(0) |
| Erythromycin | 15(57.69) | 9(34.62) | 2(3.85) |
| Clindamycin | 19(73.07) | 6(14.93) | 0(0) |
| Cloxacillin | 17(65.38) | 7(26.92) | 2(7.69) |
| Tetracycline | 5(19.23) | 7(26.92) | 14(53.85) |

## Discussion

The overall prevalence of *L. monocytogenes* from both sample types was found to be 26 (3.39%), which was characterized and confirmed both phenotypically and molecularly. The current finding is in line with the findings of [51] (2.6%), [52] (2.94%), [53] (4%), [30] (4.1%), [54] (4.3%), [55] (4.8%), [27] (5.1%), [56] (5.3%), [57] (5.4%), and [58] (6.25%). However, it is lower than the findings of [59] (12.4%), [60] (17%), [61] (26.66%), and [62] (32.7%). But it is higher than the report of [63] (1.5%).

The meat sample based prevalence of *L. monocytogenes* was found to be 4.17 %. This is in agreement with the findings of [57] (2.6%), [64] (3.3%), [53] (3.92%), [58] (6.66%). However, it is lower than the reports of [52] (8%), [65] (9.6%), [59] (10.3%), [66] (12%), [60] (14%), [67] (15%), [68] (15.4%), [51] (18.7%), and [69] (45.6%). But it is higher than the finding of [63] (0%). Likewise, the milk sample based prevalence of *L. monocytogenes* was found to be 2.6%. The current report is in agreement with the reports of [70] (0.41%), [62] (1.1%), [58] (4.0%), [71] (4%), [63] (4.0%), [72] (4.29%), [73] (5%), and [74] (5.1%). But it is lower than the reports of [75] (5.6%), [76] (7.54%), [57] (13%), [54] (5.3%), [52] (8%), [77] (8.8%), [51] (18.1%), [78] (21.7%), and [61] (40%). In general, the variation both in the overall and sample wise prevalence rates of *L. monocytogenes* might be due to differences in sample types of foods of bovine origin, sources of the samples, processing plants, approaches of sample collection, sample size, methodological approaches, isolation and identification techniques, prevalence calculation/interpretation, geographical locations, hygienic conditions, handling and transportation of samples, and contamination rates from utensils and personnel. The serovars that were identified in the current study were belonged to 1/2b and 4b. [73] and [69] were also reported these serovars in addition to some others. In general, the most common causes of human listeriosis among the 13 serotypes of *L. monocytogenes* are 1/2a, 1/2b, and 4b, and of these serotype 4b has been related to the most recent outbreaks of listeriosis, and serotypes 1/2a and 4b are commonly reported in animals [14-20, 25]. The presence of molecular serogroup 1/2b and 4b isolates (potential serotype 4b) in food of bovine origin may pose great health threat, since *L. monocytogenes* 4b has caused numerous human listeriosis outbreaks [79].

In the present study, the 26 *L. monocytogenes* isolates were investigated for their *in vitro* antimicrobial susceptibility pattern. The antimicrobial susceptibility results indicated as large proportions of the isolates were found to be highly susceptible to Ampicillin (88.46%) and Vancomycin (84.62%). However, the isolates had shown the highest level of resistance against Nalidixic Acid (96.15%). The highest intermediate was observed to Amoxicillin (57.69%). This is in agreement with the reports of [58] who reported a high degree of resistant against Nalidixic Acid (50%) and Tetracycline (37.5%) but the highest level of sensitivity to Vancomycin (100%); [60] who reported 97% of susceptibility to Ampicillin and Vancomycin, [65] who reported 100% of susceptibility Vancomycin, [61] who reported 100% of susceptibility to Ampicillin and Vancomycin, [77] who reported 100% of susceptibility to Ampicillin, Gentamycin, and Vancomycin and 24.2% of resistance against Tetracycline, [73] who reported a high level of sensitivity Ampicillin (78.9%), Penicillin G (77%), Erythromycin (71.2%), and Vancomycin (67.3%), [62] who reported 100% of susceptibility to Vancomycin and 96.4% of resistant against Nalidixic Acid. This is contradicted with the findings of [80] who reported a high degree of resistance against Amoxicillin (50%) and Vancomycin (50%), [76] who reported a high degree of resistance against Clindamycin (100%), Ampicillin (95.7%), Erythromycin (95.7%), Penicillin (91.3%), Tetracyclin (82.6%), Streptomycin (78.3%), and Vancomycin (69.7%), [58] who reported a high degree of resistance against Penicillin (66.7%) and the highest level of sensitivity to Amoxicillin (100%), Cloxacillin (100%), and Gentamicin (100%). It is similar to the findings of [59] who reported 13.6% of intermediate response to Ciprofloxacin. Moreover, 23.10%, 34.62%, and 42.31% of the isolates were showed resistance for one, two, and more than two drugs, respectively. This is lower than the findings of [76] who reported a 96.9% of multidrug resistance. But higher than the report of [58]) (16.7%), and [81] who reported a 2% of resistance to two antibiotics and 2% resistance to three antibiotics, [73] who reported that 15.3% the isolates were resistant to only one drug and 36.5% were resistant to multiple drugs. But is lower than the reports of [59] who reported 72.3% of multiple drugs resistant. In general, *Listeria monocytogenes* is usually susceptible to a wide range of antibiotics. Nevertheless, evolution of bacterial resistance towards antibiotics has been accelerated considerably by the selective pressure exerted by over prescription of drugs in clinical settings and their heavy use as promoters in animals’ husbandry [35]. Moreover, the genes responsible for antibiotic resistance could be transferred through movable genetic elements such as conjugative transposons, mobilizable plasmids, and self-transferable plasmids to other foodborne bacteria in the gastrointestinal tract. In *Listeria spp*., efflux pumps have also been reported as the resistant mechanism [82]. Hence, the use of antimicrobials in veterinary medicine is the main cause of the development of AMR foodborne bacterial pathogens including *L. monocytogenes*, as AMR pathogens can easily be transported from animal to human via food consumptions [83].

### Conclusion and Recommendations

In conclusion, the current study revealed the occurrence of *L. monocytogenes* with different pathogenic serovars (1/2b and 4b) in raw meat and milk samples of bovine origin collected from different sites of the study area and development of antimicrobial resistance to one, two, and multi to the tested antimicrobials. Hence, regardless of its prevalence rate foods of bovine origin can serve as a potential vehicle for transmitting *L. monocytogenes* and can be risk of infection to consumers of raw or under cooked milk and meat and are of significant concern for public health and give a warning signal for the possible occurrence of a foodborne outbreak in humans in the study area. It has an important public health implication that when antibiotic resistant *L. monocytogenes* is present in raw foods especially in developing countries like Ethiopia where there is very common practice of consumption of raw meat and milk products and widespread and uncontrolled use of antibiotics. As a result, both its occurrence and development of a multi-drug resistance should receive particular attention and is an alert for the concerned bodies. Thus, a coordinated effort is imperative to reduce or eliminate the risk posed by this pathogen at various stages in the food chains and on the controlled and careful use of antimicrobials both in veterinary and human treatment regimes. Moreover, awareness creation should be made on foodborne disease caused by *L. monocytogenes* with due consideration on the safe handling and consumption of foods of bovine origin and selection and safe use of antimicrobials. Finally, nationwide molecular epidemiology, phenotypic and molecular characterization, in-depth typing and drug resistant gene identification of *L. monocytogenes* should be undertaken.

## Acknowledgements

The authors would like to acknowledge owners, managers and workers of the different dairy farms, abattoir, butcher shop, restaurant, milk whole seller, and cafeteria of the study site for their keen interest and cooperation during the collection of the samples, and to the College of Veterinary Sciences staff members who directly or indirectly helped us during the research period.

